# Prevalent phenotypic mutation impairs binding of broadly neutralizing antibodies to influenza hemagglutinin

**DOI:** 10.64898/2026.07.24.740633

**Authors:** Margarita Barriga, Olga Barranco-Gómez, Laura García-Corzo, Alberto Fernandez-Fernandez, Antón Vizcaíno, Paula Ramilo, Antonio Osuna, Valeria A. Risso, Jose M. Sanchez-Ruiz

**Author notes:** Corresponding author. Contact: Jose M. Sanchez-Ruiz. Departamento de Quimica Fisica. Facultad de Ciencias, Universidad de Granada, 18071 Granada, Spain. These authors contributed equally (Margarita Barriga, Olga Barranco-Gómez, Laura García-Corzo, Alberto Fernandez-Fernandez).

## Abstract

Mistakes during protein synthesis, such as transcription errors, occur often and lead to non-inheritable amino acid replacements generally known as phenotypic mutations. We recently used a consensus approach in high-throughput sequencing to determine the error landscape for influenza-hemagglutinin mRNA. We found single-site errors to occur with widely different frequencies. Here we show that the most prevalent transcription error encodes a phenotypic mutation that impairs binding of broadly neutralizing antibodies. The error occurs in 0.2-0.5% of mRNA molecules and, consequently, many virions will expose hemagglutinin variants bearing the encoded amino acid replacement. Our results point to a mechanism of antibody evasion, akin to programmed recoding, in which evading mutations are encoded by transcription errors promoted by inheritable RNA sequence/structure patterns.

## INTRODUCTION

Mistakes during protein synthesis (translation and transcription errors) lead to amino acid replacements generally known as phenotypic mutations. While not directly inheritable, phenotypic mutations are orders of magnitude more frequent than genetic mutations and have been proposed to play various evolutionary roles (Drummond and Wilke, 2009; Goldsmith and Tawfik, 2009; Evans et al., 2018; Schmutzer and Wagner, 2023; Landerer et al., 2024). Viral burst sizes (*i*.*e*., the numbers of new virions generated through the infection of a single host cell) are typically very large (Philips and Milo, 2015). Consequently, viruses can achieve huge amplifications on the basis of a few successful biomolecular interactions (Luzon-Hidalgo et al., 2025). Viruses thus appear well prepared to take advantage of the molecular diversity generated by protein synthesis mistakes (Luzon-Hidalgo et al., 2021).

Influenza hemagglutinin is the protein responsible for the key interaction of the influenza virion with the host cell. It is also the main target for neutralizing antibodies. Recently (Barranco-Gomez et al., 2026), we used a model influenza strain (A/WSN/1933 H1N1) and a consensus approach in high-throughput sequencing to assess the single-site error landscape for the two hemagglutinin-encoding RNAs: the messenger mRNA and the viral (virion encapsulated) vRNA. We found single-site errors to occur at widely different frequencies. Such a highly non-random error landscape may determine influenza evolution to some extent (Barranco-Gomez et al., 2026). Moreover, a highly non-random error landscape suggests that mRNA biosynthesis errors, even if not inheritable themselves, can be promoted by inheritable RNA sequence/structure patterns.

Here, we focus on the most prevalent variation in the error landscape for hemagglutinin-encoding mRNA, a 920c>t substitution that is detected in about 0.2-0.5% of the mRNA molecules. 920c>t occurs at a much lower level in the viral vRNA. Therefore, its prevalence in mRNA plausibly reflects a promoted transcription error. 920c>t encodes a P307L amino acid replacement, which is intriguing. Position 307 is in the middle of the hemagglutinin stem close to epitopes targeted by broadly neutralizing antibodies. A proline to leucine replacement may be disruptive (coefficient -3 in BLOSUM 62 matrix; Henikoff and Henikoff, 1992) and cause local conformational alterations in hemagglutinin structure that could impair antibody binding. Indeed, ELISA assays and biolayer interferometry experiments reported in this work confirm that P307L impairs binding of broadly neutralizing antibodies to influenza hemagglutinin.

Our results point to a mechanism of antibody evasion, akin to programmed recoding (Brierley et al., 2010; Miller and Griedoc, 2010; Namy and Russel, 2010), in which the evading mutation is encoded by a transcription error promoted by inheritable RNA sequence/structure patterns. Note that there are up to about 1500 hemagglutinin monomers at the surface of one influenza virion (Sautto et al., 2018). A prevalence of 0.2-0.5% for the 920c>t variation in mRNA implies that most virions will expose hemagglutinin variants bearing the amino acid replacement that could enable evasion of broadly neutralizing antibodies. Moreover, the stochastic nature of mRNA synthesis and translation in single host cells makes it plausible that some virions harbour comparatively large numbers of the hemagglutinin variants. Further note that, given the large burst sizes for influenza (between about 1000 and 10000 virions generated through the infection of single mammalian cell: Philips and Milo, 2015), virus propagation could be sustained even if only a comparatively small fraction of the virions are competent for infection.

Unlike the highly variable head region, the stem of influenza hemagglutinin is largely conserved, plausibly reflecting functional constraints. Antibodies targeting the stem region are of considerable interest because they could provide the basis for broad protection against various strains of the virus. The P307L replacement, however, could enable the virus to evade broadly neutralizing antibodies. The fact that the replacement is encoded as a promoted phenotypic mutation, rather than as directly inheritable genetic mutation, may serve as a mechanism to bypass functional trade-offs and to avoid detection by the immune system (see Discussion).

## RESULTS

In our recent work (Barranco-Gomez et al., 2026), the error landscape for hemagglutinin-encoding mRNA was determined for two completely independent biological samples, mRNA-(1) and mRNA-(2). An excellent agreement was found between data from the two samples, in particular for the high frequency errors. The most prevalent biosynthesis error in both mRNA replicas is 920c>t (Figure 1A). We also studied (Barranco-Gomez et al., 2026) the error landscape for two independent samples of the viral (*i*.*e*., virion-encapsulated) vRNA. As shown in Table 1, the frequency of the 920c>t variation is substantially lower in the vRNA samples (about 1.4·10^-4^) as compared with the mRNA samples (2-5·10^-3^). Both vRNA and mRNA are synthesized by the viral RNA-dependent RNA-polymerase (Bolvin et al., 2010; Fan et al., 2019). vRNA is replicated in two stages through an intermediate cRNA copy, while mRNA is being transcribed in a single stage from a vRNA template. The higher frequencies in the mRNA samples as compared with the vRNA samples thus support that the 920c>t variation is mostly generated as a transcription error in the synthesis of mRNA from vRNA.

**Table 1.** Determination of the frequency of the 920c>t variation that encodes for the P307L amino acid replacement.

| Biological sample | Total depth | Reference allele (C) depth | Alternative allele (T) depth | Frequency |
| --- | --- | --- | --- | --- |
| mRNA-(1) | 787887 | 786081 | 1577 | $2.0 \cdot 10^{-3}$ |
| mRNA-(2) | 614802 | 611733 | 3061 | $5.0 \cdot 10^{-3}$ |
| vRNA-(1) | 65020 | 65008 | 9 | $1.4 \cdot 10^{-4}$ |
| vRNA-(2) | 27454 | 27450 | 4 | $1.5 \cdot 10^{-4}$ |

**Figure 1.**
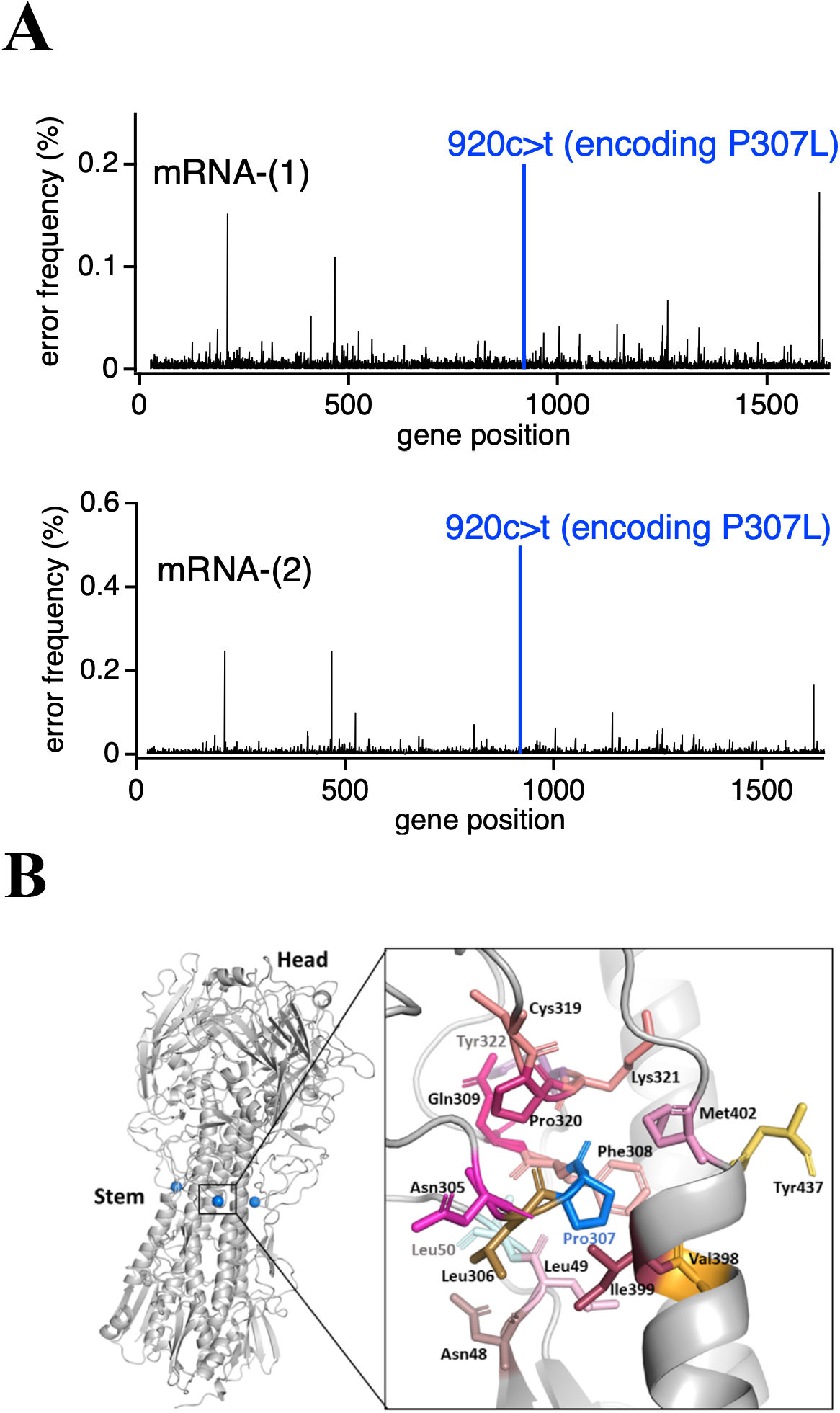
Most prevalent variation in the error landscape for hemagglutinin-encoding mRNA and molecular environment of proline at position 307. (A) Biosynthesis error landscape for hemagglutinin-encoding mRNA (data from Barranco-Gomez et al., 2026). Two independent biological samples, mRNA-(1) and mRNA-(2), were studied. Data are shown as plots of error frequency versus gene position. As previously discussed (Barranco-Gomez et al., 2026), there is good congruence between the two sets of data. The most prevalent variation on both replicas is 920c>t which encodes for the P307L amino acid replacement. (B) Molecular environment of the proline at position 307 in the 3D-structure of influenza hemagglutinin. The inset highlights the amino acid residues close to P307, as identified by a distance between closest atoms smaller than 6 Amstrong. The hemagglutinin 3D-structure was generated by SWISS-MODEL using as template the X-ray structure of PDB ID 6N41, which corresponds to a hemagglutinin with 88% sequence identity with the hemagglutinin from A/WSN/1933 used in this work.

920c>t encodes for a P307L amino acid replacement. Figure 1B shows the location of position 307 in the 3D-structure of hemagglutinin. Proline at position 307 is largely buried (solvent accessibility of around 5%) and interacts with a diversity of amino acid residues (see the magnified inset in Figure 1B). Proline and leucine are quite different in terms of interaction energetics with other residues and in terms of conformational preferences.

Replacement of a buried and well-packed proline with a leucine residue will therefore cause substantial structural perturbation and local conformational rearrangements. Furthermore, position 307 is in the middle of the stem, a region of the hemagglutinin structure that provides epitopes for broadly neutralizing antibodies (Figure 2).

**Figure 2.**
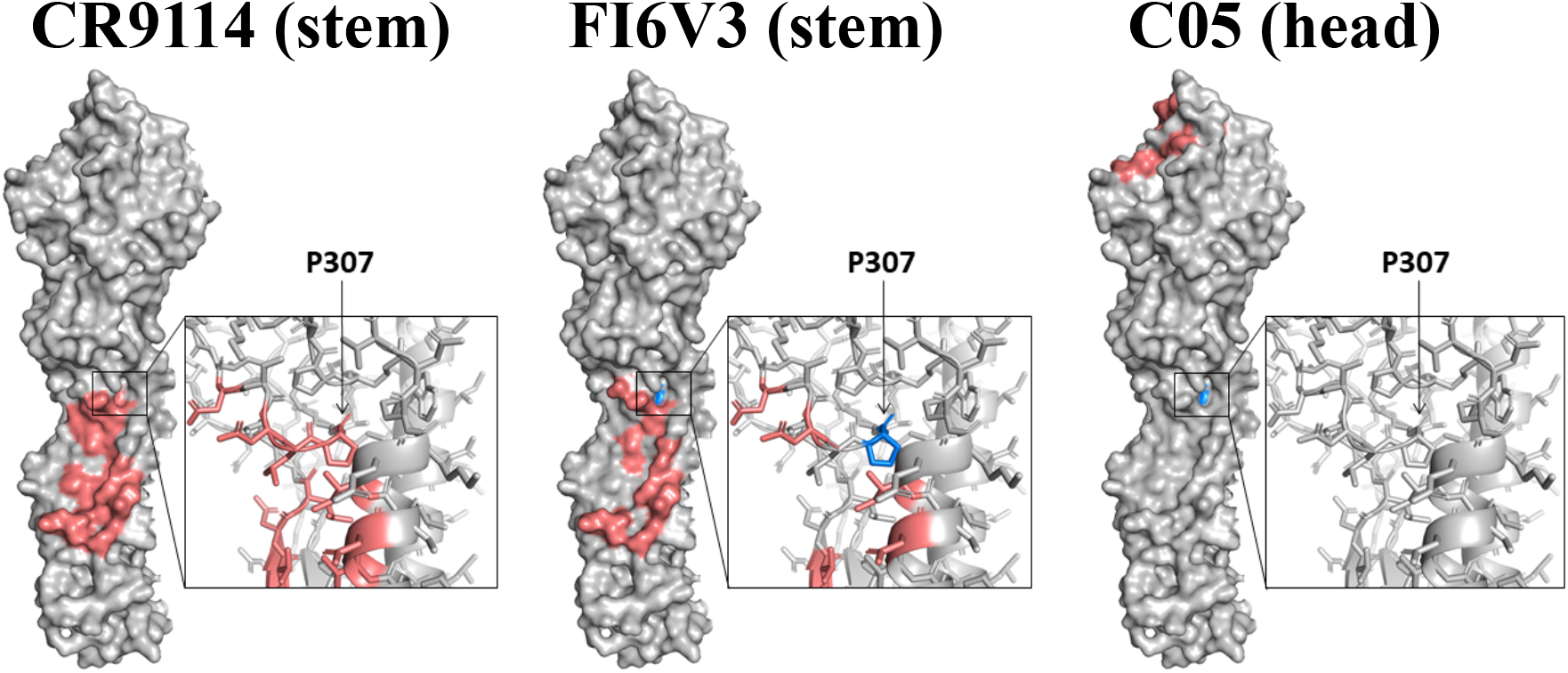
Structural relation of proline at position 307 with the epitopes for the several antibodies studied in this work. Position 307 is at or close to the epitopes for the broadly neutralizing antibodies that target the stem (CR9114 and FI6V3), while it is far from the epitope for the C05 antibody that targets the head. The hemagglutinin 3D-structure was generated as explained in the legend to figure 1 and the location of the epitopes was derived from the literature (Corti et al., 2011; Dreyfus et al., 2012; Ekiert et al., 2012).

Overall, the possibility arises that the P307L amino acid replacement impairs binding of broadly neutralizing antibodies by disrupting hemagglutinin structure in the middle of the stem. To test this possibility, we prepared wild-type hemagglutinin and its P307L variant through expression in human HEK293 cells (see Methods for details), and we studied their interaction with the broadly neutralizing antibodies CR9114 (Dreyfus et al., 2012) and FI6v3 (Corti et al., 2011). As a control, we also studied the antibody C05 (Ekyert et al., 2012) which does not target the stem but binds to an epitope in the head of the hemagglutinin molecule.

The epitopes for the three antibodies (Corti et al., 2011; Dreyfus et al., 2012; Ekyert et al., 2012; Dzimianski et al., 2023) are shown in Figure 2. Position 307 is part of (or very close to) the epitopes for the stem-antibodies (CR9114 and FI6v3) and far from the epitope for the head-antibody (C05). Note that we prepared hemagglutinin lacking the 529-565 residues which corresponds to the transmembrane domain. That is, we used the hemagglutinin ectodomains (extracellular domains) in all the interaction experiments described below.

We first studied the hemagglutinin-antibody interactions using ELISA assays (Figure 3). The profiles for the interaction of the stem-antibodies with wild-type hemagglutinin appear sigmoidal and indicate binding with nanomolar or subnanomolar affinity. On the other hand, the profiles for the interaction of the stem-antibodies with the variant are not sigmoidal, but typically show weak baseline-like dependencies, supporting that antibody binding has been substantially impaired by the P307L mutation. This conclusion is robust, as we performed several independent experiments for each antibody using different experimental protocols and conditions (Figure 3, Table 2 and section entitled “Robustness and reproducibility of ELISA assays” in Methods). ELISA experiments with the control C05 head-antibody reveal a tight binding which is not noticeably impaired by the P307L replacement (Figure 2). Therefore, impairment of the binding of the stem-antibodies cannot be put down to a global unfolding (or misfolding) of the hemagglutinin caused by the P307L replacement. Rather, the impairment is likely linked to local conformational re-arrangements in the stem that do not have a major effect on the head region.

**Table 2.** Assessing the robustness and reproducibility of ELISA assays.

| Antibody | Figure | Panel | Temperature (°C) | Incubation time | Washing Machine | Experimental Team |
| --- | --- | --- | --- | --- | --- | --- |
| **C05** | 3 | A1 | 4 | overnight | WM2 | 1 |
|  | 3 | A2 | 4 | overnight | WM2 | 1 |
|  | 3 | A3 | 25 | 1 hour | WM3 | 1 |
| **CR9114** | 3 | B1 | 4 | overnight | WM2 | 1 |
|  | 3 | B2 | 4 | overnight | WM2 | 1 |
|  | 3 | B3 | 4 | overnight | WM3 | 1 |
|  | 3 | B4 | 25 | 1 hour | WM3 | 1 |
| **FI6v3** | 3 | C1 | 4 | overnight | WM1 | 2 |
|  | 3 | C2 | 4 | overnight | WM3 | 2 |
|  | 3 | C3 | 4 | overnight | WM3 | 2 |
|  | 3 | C4 | 25 | 1 hour | WM3 | 1 |

**Figure 3.**
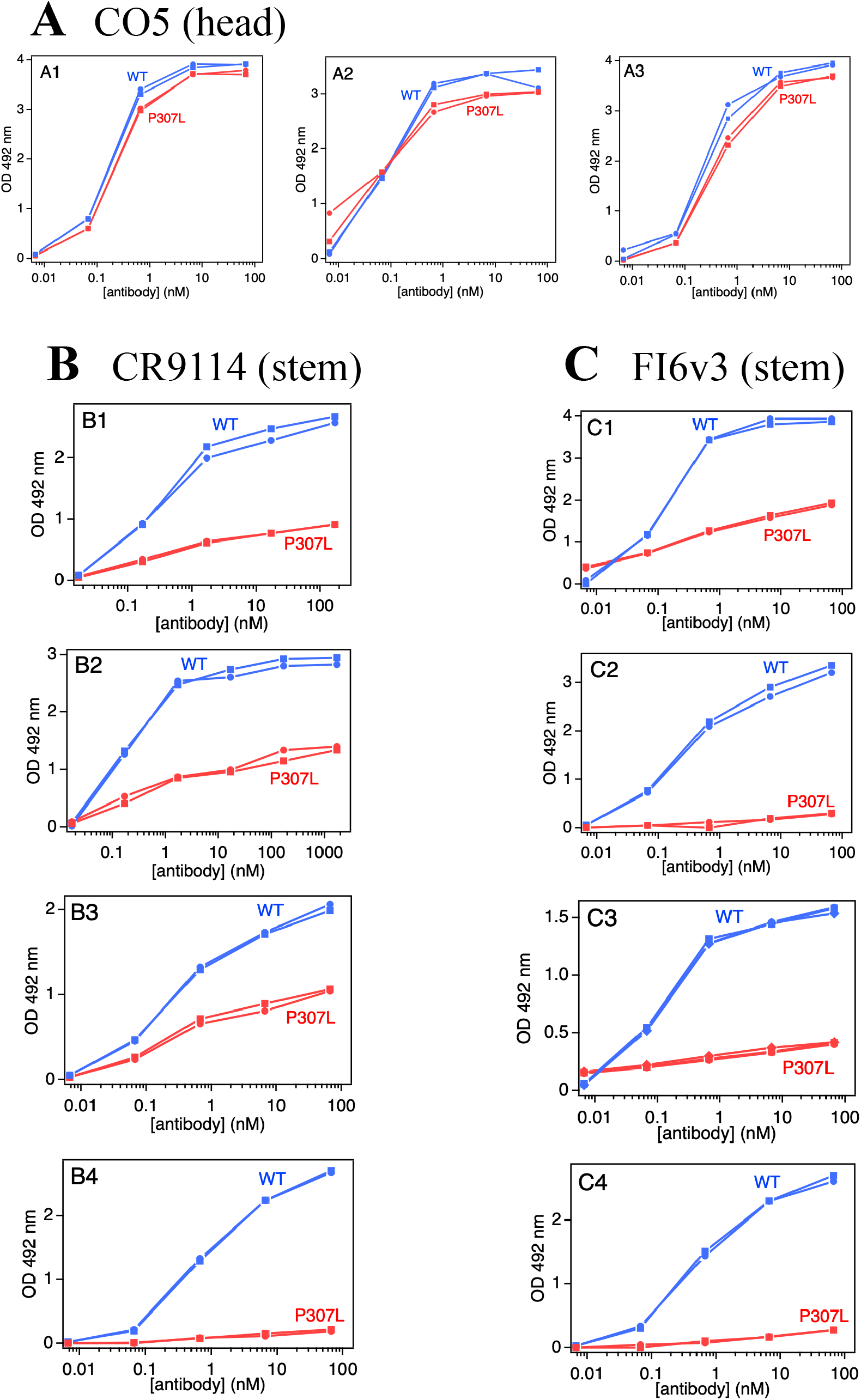
ELISA assays of the interaction of several antibodies with the wild-type form and P307L variant of influenza hemagglutinin. For each assay, two or three measurements were taken (shown with different data symbols). Several independent replica assays were performed for each antibody-hemagglutinin couple (given in separate panels). These replicas involved different researchers, different washers and different experimental conditions (see Table 2 and section entitled “Robustness and reproducibility of ELISA assays” in Methods).

We further explored the antibody-hemagglutinin interactions using biolayer interferometry (BLI). In these experiments (see Methods for details), the antibody was immobilized onto the sensor and, to obtain adequate signal, we used hemagglutinin concentrations that were substantially higher than the affinities (nanomolar to subnanomolar) derived from ELISA assays for the antibody interactions with the wild-type hemagglutinin. Figure 4A shows BLI profiles obtained at 25 ºC with a hemagglutinin ectodomains concentration of 79 nM. At this very high hemagglutinin concentration, interaction is detected in all cases. Yet, binding of the stem-antibodies (CR9114 and FI6v3) to the P307L variant of hemagglutinin is clearly impaired, as shown by the smaller amplitudes of the binding profiles. In all cases, the dissociation profiles display extremely slow kinetics, from which we could not derive reliable values for the dissociation rate constant, k_OFF_. Therefore, we did not attempt to derive estimates of binding affinities from k_ON_/k_OFF_ ratios. Interestingly, the experiments shown in Figure 3A also seem to suggest that binding kinetics is slower for the P307L variant, as compared with the wild-type form. We therefore performed an additional set of experiments (Figure 4B) at 37 ºC, using comparatively low sensor loadings, a lower hemagglutinin concentration of 26.3 nM and longer measurement times. These conditions were selected with the aim of facilitating a more reliable assessment of slow binding kinetics. The experiments shown in Figure 3B confirm the impaired binding of the stem-antibodies to the P307 variant. Slower binding kinetics is also visually apparent (see also the estimates of the relaxation time values in Figure 3B).

**Figure 4.**
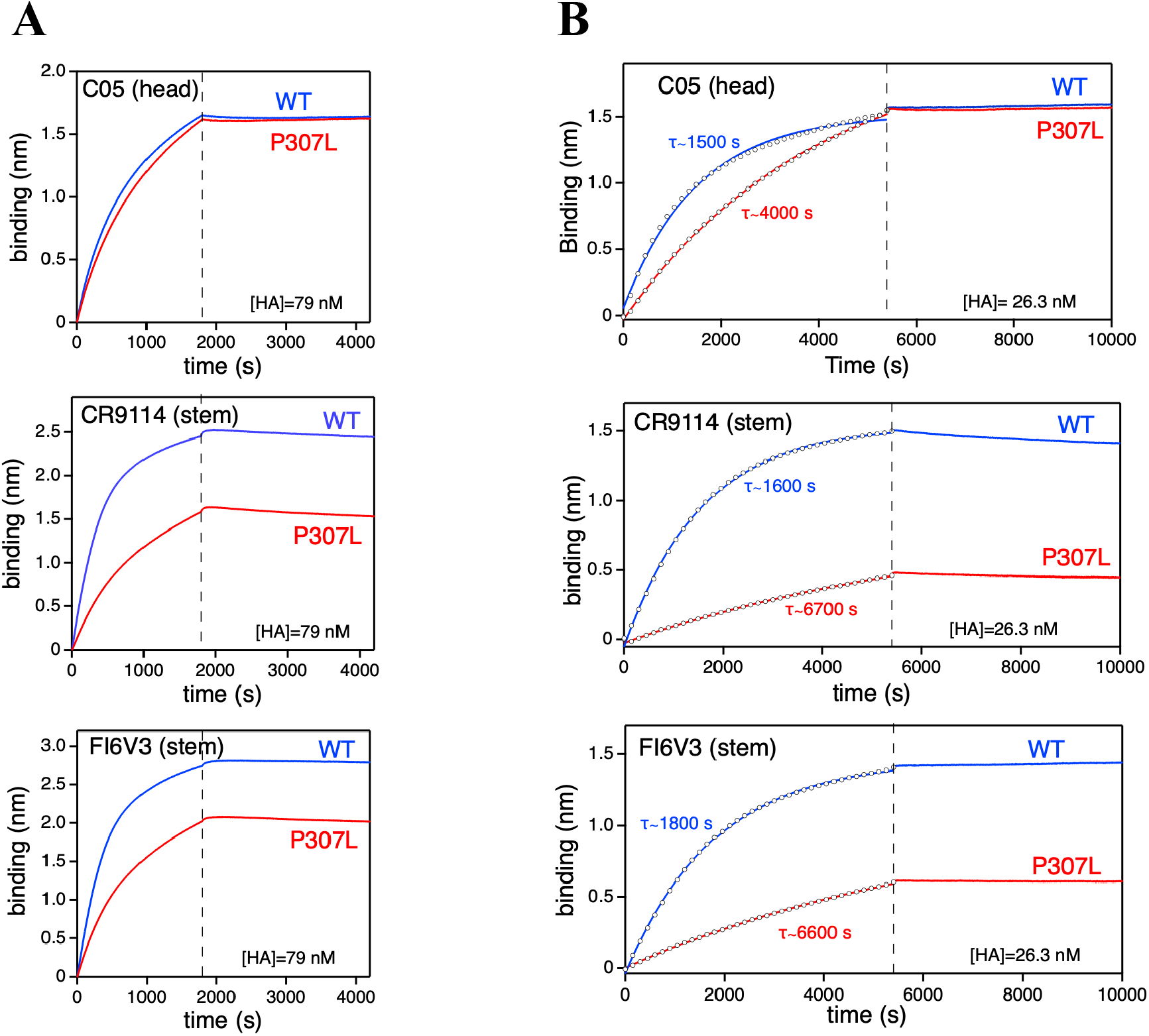
Biolayer interferometry (BLI) experiments on the interaction of several antibodies with the wild-type form and the P307L variant of influenza hemagglutinin. Antibodies were immobilized onto the sensor and exposed to the high concentrations of hemagglutinin ectodomains (HA). The panels display the binding and the dissociation profiles (at the left and the right of the vertical dashed line, respectively). (A) BLI experiments performed at 25 ºC, with a hemagglutinin concentration of 79 nm and using a comparatively high sensor load (antibody concentration in the loading solution of 10 μg/mL). (B) BLI experiments performed at 37 ºC, with a hemagglutinin concentration of 26.3 nM and using a low sensor load (antibody concentration in the loading solution of 1.25 μg/mL). Experimental data for the binding profiles are shown by circles. The estimates of the binding relaxation times shown were obtained from the fits of single exponentials to experimental binding data. The fits are shown as continuous lines. To facilitate the visual comparison of the fitted dependencies with the experimental binding data, the latter are shown as circles.

The results of the BLI experiments seem consistent with the impairment of the binding of stem-targeting antibodies (CR9114 and FI6v3) being due to the disruptive nature of the P307L mutation and its consequent impact on local conformational diversity. That is, replacement of proline at position 307 with leucine should disrupt stabilizing interactions thus generating an ensemble of structures that differ mostly in the local conformation around position 307. Most of the conformations in the structural ensemble will be unable to bind the stem-antibodies, but a few conformations could still be competent for antibody binding, accounting for the much-diminished overall affinity and the slower binding kinetics. On the other hand, once the antibody is bound to a competent conformation, the hemagglutinin-antibody intermolecular interactions will be strong and the dissociation kinetics will be slow, as the experiments show.

## DISCUSSION

Unlike the highly variable head region, the stem of influenza hemagglutinin is largely conserved. Antibodies targeting the stem region are thus of considerable interest because they could provide the basis for broad protection against various strains of the virus. However, we have shown that the most prevalent error in the biosynthesis of influenza hemagglutinin mRNA encodes an amino acid replacement that impairs binding of broadly neutralizing stem-antibodies. We have derived this result from studies on hemagglutinin and hemagglutinin-encoding mRNA from influenza virus strain A/WSN/1933 H1N1. Yet, it is plausible that the capability of the P307L mutation to impair binding to broadly neutralizing antibodies also applies to other strains, given the conservation of the stem region in general and, specifically, the high conservation of proline at position 307 in hemagglutinin. A search in the VarEPS-Influ:an database (Shu et al., 2023) shows that, out of about 9·10^5^ sequences of H1 hemagglutinin reported in this database, only about 10 do not bear proline at position 307.

Errors in the synthesis of mRNA are not inherited. Therefore, our results point to a mechanism of antibody evasion in which the evading mutations are encoded by transcription errors promoted by inheritable RNA sequence/structure patterns. This mechanism would be akin to programmed recoding, which is used by some viruses to generate functional viral proteins at significant levels (Brierley et al., 2010; Miller and Griedoc, 2010; Namy and Russel, 2010). In programmed recoding, sequence/structure motifs in the viral genetic material promote events, such as stop-codon readthrough and frameshifting, that would normally be considered as gene-expression errors. The plausibility of this kind of mechanism in the context of antibody evasion by the influenza virus is elaborated below.

There are up to about 500 hemagglutinin trimers on the surface of a single influenza virion (Sautto et al., 2018), which means that there are up to about 1500 hemagglutinin monomers per virion. According to our sequencing data (Figure 1A), the error that encodes the P307L mutation is present in about 0.2-0.5% of the molecules of hemagglutinin mRNA. Therefore, the surface of a single virion is expected to expose up to about 3-7 hemagglutinin monomers bearing the P307L mutation as a result of transcription errors. This is, of course, an estimated average. In fact, the stochastic nature of mRNA synthesis and translation in single host cells makes it plausible that some virions harbour comparatively large numbers of hemagglutinin variants. Note that there is a wide variability in mRNA levels between single host cells (Russel et al., 2018) and that a single mRNA molecule is translated many times (Traverse and Ochman, 2016). Therefore, a transcription error that occurs when there are comparatively few viral mRNA molecules present per cell will likely generate virions with much larger numbers of hemagglutinin variants at the surface and highly competent for infection in the presence of neutralizing antibodies. It is also important to note that virus propagation could be sustained even if only a comparatively small fraction of the virions infect host cells, because the influenza burst size (the number of virions generated through the infection of single host cell) is enormous, with values between 1000 and 10000 virions reported for mammalian cells (Philips and Milo, 2015).

Finally, we may ask why evolution would have selected to encode antibody evasion as a promoted transcription error in the synthesis of mRNA rather than as a directly inheritable genetic mutation in the vRNA. One possibility is that the P307L amino acid replacement involves a trade-off between escape from antibodies and functionality. That is, the local conformational change brought about by the disruptive P307L replacement may enable escape from broadly neutralizing antibodies but at the cost of impairing to some extent the primary function of the stem as mediator of viral entry. Therefore, the presence of the substitution that encodes P307L at the genetic level would make virions less competent for infection in the absence of the neutralizing antibodies. In fact, the level of the 920c>t error in the viral vRNA, which is subject to selection, is about 0.01%, substantially lower than the corresponding error in mRNA (Table 1). On the other hand, encoding P307L as a promoted transcription error would guarantee that the virus still propagates after antibodies targeting the stem have been generated, while allowing efficient propagation prior to antibody generation. Also encoding as a promoted transcription error would ensure that the variant bearing the evading mutation is always expressed at low level, which may help avoid its detection by the immune system.

## Acknowledgments

This work was supported by Grant IHRC22/00004 (to J.M.S.-R.) funded by the “Instituto de Salud Carlos III” and Next Generation EU.

## Competing interests

The authors declare no competing interests

## Data availability

All relevant data are provided in Figures 1, 3 and 4. Tables including these data are available from the authors upon request. Error frequency data are provided in in our recent work (Barranco-Gomez et al., 2026).

## METHODS

### Single-site error landscape for hemagglutinin-encoding RNA

The biosynthesis error landscapes for hemagglutinin-encoding mRNAs analyzed here (Table 1 and Figure 1) are taken from our recent work (Barranco-Gomez et al., 2026) in which we used a model influenza strain (A/WSN/1933 H1N1) and a unique-molecular-identifier-based high throughput sequencing approach to assess the single-site error frequencies.

### Preparation of hemagglutinin ectodomains

Hemagglutinin from the A/WSN/1933 H1N1 strain was used in ELISA and BLI experiments. Two different recombinant hemagglutinin proteins were used: a wildtype (WT) and a P307L variant (H1 numbering, starting from the initial methionine). In both cases, the ectodomains (*i*.*e*., the extracellular domains lacking the transmembrane domain) were used. Specifically, Met1 to Gln528 proteins (*i*.*e*., proteins lacking the 529-565 residues that correspond to the transmembrane domain) were used with a polyhistidine tag attached to the C-terminal for purification. The recombinant proteins were prepared by the Custom Service of Sino Biological using the protocols and quality controls they routinely employ to produce recombinant hemagglutinin proteins in Human Embryonic Kidney 293 Cells (HEK293). The proteins were purified by affinity chromatography, and their purity was checked by SDS-PAGE. Size Exclusion Chromatography coupled with High-Performance Liquid Chromatography (SEC-HPLC) indicated that the proteins were 80% trimeric and 20% monomeric. Aqueous solutions of the proteins in PBS (phosphate buffered saline solution) were shipped to our lab in dry ice.

### Enzyme-linked immunosorbent assays (ELISA)

ELISA assays were used to assess the interaction of several neutralizing antibodies with the WT and P307L variants of the hemagglutinin ectodomains from influenza strain A/WSN/1933 H1N1. The antibodies studied were CR9114 (stem) (human, DVV03807 AntibodySystem), FI6V3 (stem) (human, PAB-214 Creative Biolabs) or C05 (head) (human, PABL223 Creative Biolabs). Specific experimental details are given below:

Stock solutions (0.01 μg/μL) of either the WT ectodomain or the P307L variant of the ectodomain were diluted in PBS (phosphate buffered saline solution), pH 7.4 (Gibco, 10010-031). 100-μl aliquots of this solution were placed into the wells of a 96-well ELISA Medisorp microplate (Thermo Scientific, 467320). After overnight incubation at 4 °C, plates were washed four times with washing buffer: PBS containing 0.05% Tween 20 (P9416 Sigma Aldrich). After washing, 300 μl of blocking buffer (5% non-fat milk in washing buffer) was added to each well. After incubation for 3 hours at 25 °C, 100 μl of antibody solutions in blocking buffer were added to the wells. Serial dilutions 1/10 were performed, and each different condition was tested in duplicate or triplicate. Overnight incubations at 4 °C or 1-hour incubations at 25 ºC were performed. After washing four times with washing buffer, 100 μl of HRP-conjugated antibody (Mouse Anti-Human IgG Fc Antibody-50B4A9, Genscript A01854) diluted (1/5000) in PBS containing 0.05% Tween 20 was applied to each well for 1 hour at 25 °C. After washing four times with washing buffer, binding was quantified by using 100 μl of horse radish peroxidase (HRP) substrate solution, using an o-phenylenediamine (OPD) chromogenic substrate system (P8287 Sigma-Aldrich). The substrate was prepared in a citrate–phosphate buffer system (0.02M citrate, 0.05 M phosphate) supplemented with 0.02 % hydrogen peroxide (H2O2) (H1009 Sigma Aldrich). The reaction was allowed to proceed for 30 minutes, at 25 ºC, in dark conditions, and was stopped adding 50 μl of a HCl 3M solution to stabilize the color. The absorbance was then measured spectrophotometrically at 492 nm in an automated plate reader (NanoQuant infinite M200 Pro, Tecan).

### Robustness and reproducibility of ELISA assays

To assess the robustness and reproducibility of the ELISA assays, antibody binding was evaluated under multiple independent experimental configurations that differed in antibody incubation temperature, automated plate washer (WM1, WMs and WM3), and experimental team. The ELISA data presented in Figure 3 were obtained under these different experimental conditions, providing an opportunity to assess the reproducibility of the observed antibody-binding profiles across independent assays. For C05, panels A1 and A2 correspond to independent experiments performed following overnight antibody incubation at 4 °C using the WM2 plate washer, whereas A3 was performed after a 1 h antibody incubation at 25 °C using WM3; all three experiments were carried out by independent experimental team 1 (two researchers). For CR9114, panels B1 and B2 represent independent experiments performed at 4 °C using WM2, B3 was carried out at 4 °C using WM3, and B4 was performed after a 1 h antibody incubation at 25 °C using WM3; all four experiments were conducted by independent experimental team 1. For FI6V3, panel C1 corresponds to an experiment performed at 4 °C using WM1, whereas panels C2 and C3 represent independent experiments carried out at 4 °C using WM3; these three experiments were performed by independent experimental team 2 (two researchers). Panel C4 was performed after a 1 h antibody incubation at 25 °C using WM3 by independent research team 1. The conditions of the different ELISA assays are summarized in Table 2.

Overall, these experiments comprise independent ELISA assays performed using three automated plate washers (WM1–WM3), two independent experimental teams (four researchers in total), and two antibody incubation protocols (overnight at 4 °C or 1 h at 25°C), all of which produced highly consistent binding profiles, demonstrating the high reproducibility of the assay. The three plate washers represented commercially available standard ELISA plate washers from different manufacturers and located in independent laboratories, further demonstrating that the observed antibody-binding profiles were robust across different laboratory settings and commonly used ELISA instrumentation.

### Biolayer interferometry (BLI)

Binding of antibodies (CR9114, FI6v3 and C05) to ectodomains of hemagglutinin from A/WSN/1933 (both WT and the P307L variant) was studied using an Octet R4 instrument (Sartorius). Antibodies were diluted to final concentrations of 1.25 µg/mL or 10 µg/mL (depending on the experiment) in 1x Kinetics Buffer, prepared by a 1:10 dilution of Octet Kinetics Buffer 10× (Ref. 18-1105, Sartorius) in PBS pH 7.4 (Gibco, 10010-031). Antibodies were immobilized onto an antihuman IgG-Fc-coated biosensor (AHC2 Ref. 18-5142, Sartorius) and exposed to hemagglutinin ectodomain solutions PBS pH 7.4 (Gibco, 10010-031). A typical BLI assay involved six sequential steps: 1. Baseline acquisition (60 s), 2. Antibody loading onto biosensor surface (900 s) 3. Washing step (180 s) 4. Second baseline acquisition (180 s) 5. Association phase with hemagglutinin ectodomain (∼1800 or ∼5400 s depending of the experiment) 6. Dissociation phase (∼2400 or ∼4500 s depending of the experiment). Baseline, washing, and dissociation steps were performed in 1x Kinetics Buffer.

## Notes

### Competing Interest Statement

The authors have declared no competing interest.

